# SAVE: A secure cloud-based pipeline for CRISPR pooled screen deconvolution

**DOI:** 10.1101/110262

**Authors:** Hyun-Hwan Jeong, Seon Young Kim, Maxime WC Rosseaux, Huda Y Zoghbi, Zhandong Liu

## Abstract

We present a user-friendly, cloud-based, data analysis pipeline for the deconvolution of pooled screening data. This tool, termed SAVE for Screening Analysis Visual Explorer, serves a dual purpose of extracting, clustering and analyzing raw next generation sequencing files derived from pooled screening experiments while at the same time presenting them in a user-friendly way on a secure web-based platform. Moreover, SAVE serves as a useful web-based analysis pipeline for reanalysis of pooled CRISPR screening datasets. Taken together, the framework described in this study is expected to accelerate development of web-based bioinformatics tool for handling all studies which include next generation sequencing data. SAVE is available at http://save.nrihub.org.

## Introduction

Genetic screening allows for the unbiased interrogation of a genome to ask targeted questions [1]. While originally centered around random mutagenesis or arrayed RNAi [2, 3], a growing amount of studies are moving towards large-scaled, pooled approaches [4–7]. Indeed, the advent of CRISPR has heralded a dramatic increase in the number of pooled screens [8]. For instance, in the past year, the number of datasets for CRISPR screens in Gene Expression Omnibus (https://www.ncbi.nlm.nih.gov/gds) have more than tripled from the previous year (39 datasets in 2015 and 121 datasets in 2016). In these, a collection of modifiers (e.g. shRNAs or sgRNAs) are introduced to a population (e.g. in cell culture or into an organism) blindly. Following phenotypic (e.g. enriching for surviving cells following a toxic insult) or enrichment-based (e.g. FACS sorting a given fluorophore) screening, genomic DNA is extracted and enrichment of each modifier is assessed using microarray or, more commonly, next generation sequencing. Once a daunting and expensive approach to generate hypotheses in an unbiased fashion, pooled screening approaches are now more feasible and accessible than ever. Notably, cheaper sequencing costs [9], readily available libraries via non-profit entities (e.g. Addgene) and more robust gene-disruption strategies (e.g. CRISPR and CRISPRi ([10] and [11] now allow for global gene disruption-based screens in an unprecedented manner.

Nevertheless, the bioinformatics hurdle of data deconvolution following sequencing remains a roadblock for investigators with little-to-no computational knowledge. Previous tools have been generated to analyze high-throughput screening datasets but, like most bioinformatics tools, these did not always provide a good user experience (Table 1). There are several reasons to explain these shortcomings. First, these tools often require a self-installation step to use and often require the combination of several packages to work adequately. Second, raw sequence data handling is not provided and therefore users must independently manipulate the data (leading to undue errors) by uploading their data to platforms such as RIGER and ATARI [12, 13]. Third, most available tools are specialized for dropout studies such that enrichment-based studies (or any other study with more complex data involving 3 or more groups) does not fit in the analysis pipeline. Lastly, the inflexibility in statistical analysis output in these programs may lead to improper conclusions.

**Table 1.** A comparison of SAVE with existing method for the multifunctionality

| Feature | RIGER | ATARIS | MAGeCK-VISPR | ScreenProcessing | caRpools | SAVE |
| --- | --- | --- | --- | --- | --- | --- |
| Web interface | ✓ | ✓ |  |  |  | ✓ |
| Cloud service |  |  |  |  |  | ✓ |
| FASTQ file handling |  |  | ✓ | ✓ | ✓ | ✓ |
| GUI |  | ✓ |  |  | ✓ | ✓ |
| Multiple group studies |  |  | ✓ | ✓ | ✓ | ✓ |
| Paired data analysis |  |  |  |  |  | ✓ |
| Quality control |  |  | ✓ |  | ✓ | ✓ |
| Reference | [12] | [13] | [19] | [20] | [21] |  |

To overcome these issues, we generated a web-based analysis pipeline for pooled screens: SAVE. SAVE (Screening Analysis Visual Explorer, http://save.nrihub.org) is a user-friendly pipeline that lends itself to customization according to the screen at hand. In this system, the user can upload raw sequencing files confidentially, can extract valuable screening information through customizable statistical analysis and can output various end-point results set by the user.

## Methods

To generate this platform, we first had to overcome the following issues: File size and confidentiality as well as multifunctionality in output formats. First, the file size of next generation sequencing datasets from pooled screening is big (about 10GB for each sample), so it would take much time to upload every dataset directly without any manipulation. Second, given that users are uploading raw data to the server, it is critical to minimize the information uploaded for analysis, to avoid unwanted leaking of user’s data. Third, we wanted to develop a web-based bioinformatics tool that was easy to handle and generate comprehensive visualizations for those performing the screens who are not necessarily well versed in bioinformatics.

Fortunately, the client-side web technology like JavaScript and HTML5 have rapidly developed in the recent year, and those technologies have enabled the high-performance computation for large sized file handling using modern web browsers such as Google Chrome, Mozilla Firefox, and Microsoft Edge [14]. Thus, we can perform the initial steps of data analysis at the *client-side* with the technologies, allowing for rapid and safe data minimization. Since a web-based analysis system with a centralized server may run slower than the expectation of the user, one must overcome this limitation to allow for proper web-based integration. Cloud computing offers such a solution which provides powerful and scalable computational resources, such as CPU and storage, to meet users’ expected running time at an economical efficient manner.

To address the aforementioned problems, we developed SAVE (Screening Analysis Visual Explorer) - a secure cloud-based pipeline analysis and data management for high-throughput pooled CRISPR screening - and have started to provide the public service (Figure 1). This service gives a user-friendly graphical interface (Figure 2) and it minimizes the data sent to the server by preprocessing FASTQ files on the user’s computer. The only data the server gets is the counts matrix. Also, SAVE provides a scalable service through the infrastructure provided by Amazon Elastic Compute Cloud (EC2) and Amazon Simple Storage Service (S3). It supports two different types of study (dropout and enrichment based experiment) with various proper statistics (a Student’s t-test with log-transformation of counts [15], DESeq2 [16], and the inverted beta-binomial test -- a test used for paired datasets [17]). Also, users can get a visualized web report for their data and export the results via an address-encrypted link (Figure 3).

**Figure 1.**
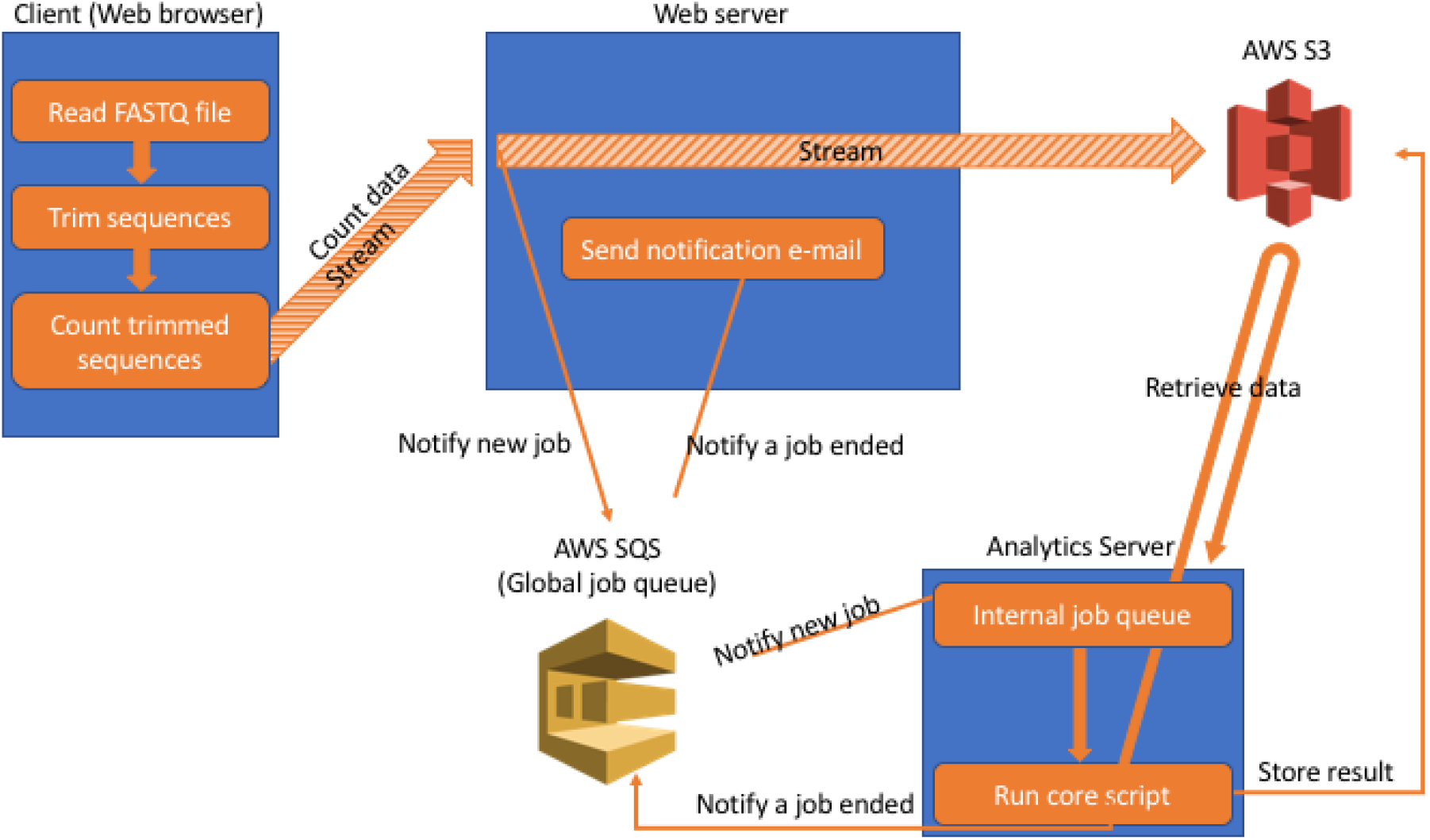
A detailed illustration of the pipeline of SAVE

**Figure 2.**
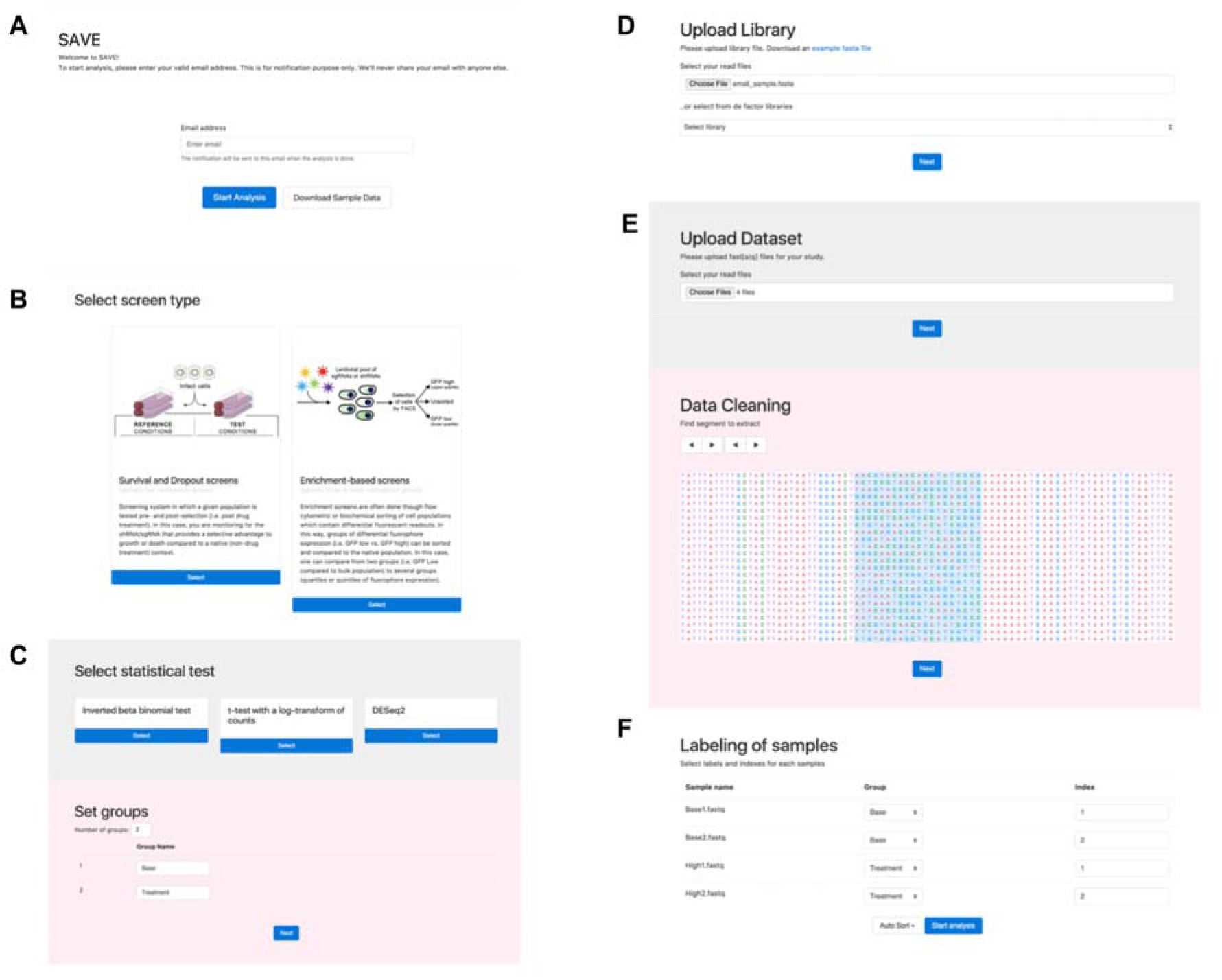
Screenshots of GUI to input user data in the SAVE website: (A) The first page to input user email address. (B) A page for the selection of a type of the experiment. (C) choice of statistical test and add labels for each group. (D) A page for the upload sgRNA Library. (E) A page for the upload FASTQ file and page to choose a region for each read of deep sequencing data to extract the sgRNA for the input sequences. (F) a page for labeling of samples to design the study.

**Figure 3.**
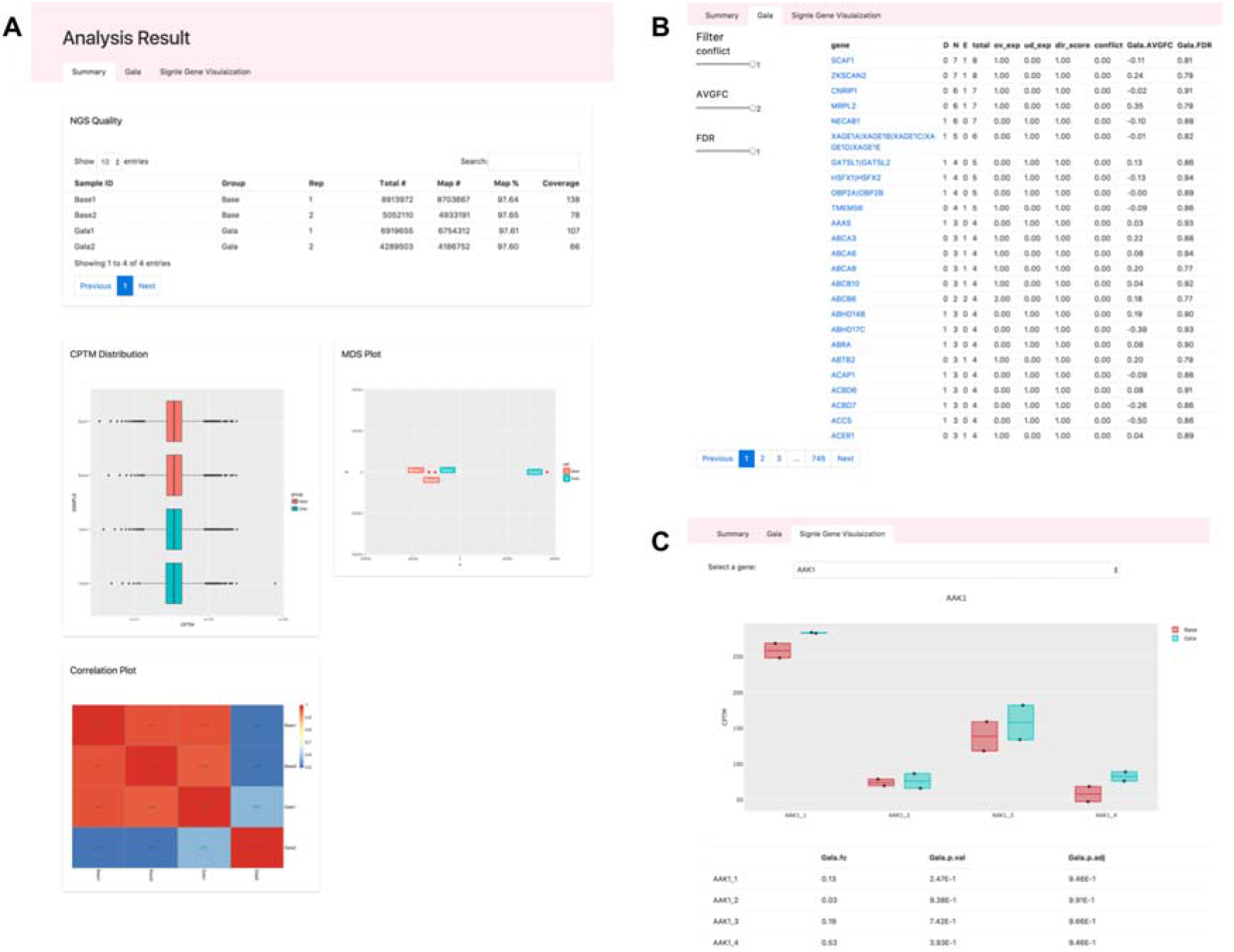
An example of the visualized report of SAVE: (A) Overview of quality of user’s deep sequencing data. (B) Statistics page to select hit genes (C) A boxplot of read count of sgRNAs for a gene

## Discussion

The decreasing costs and time-frames associated with pooled genome-wide screening approaches is incentivizing an increasing number of investigators. Nevertheless, the sequencing-to-data gap remains discouraging. We therefore developed SAVE as a solution to this issue. The multifunctionality of SAVE is clear with current established technologies such as shRNA and CRISPR but can easily be extended to any technology where short sequence barcoding is used (e.g. ORF screens and CRISPRi/a) [10] [11] [18]. Moreover, this user-friendly system allows for re-analysis of other established datasets in the context of one’s statistical analysis parameters (i.e. if you filter using certain cut-offs and want to confirm with someone else’s dataset, you can do so with SAVE). Lastly, the implications of this software extent passed screening approaches and are potentially applicable to any datasets concerning multiple sequencing files that require rapid deconvolution. One example of these would be single-cell RNA-seq experiments where massive amounts of sequencing results must be aligned, quantified and presented in a user-friendly manner.

Taken together, this study highlights a web-based platform upon which investigators can securely deposit raw sequencing files, extract the valuable data and perform statistical analyses on them, generate and prioritize hit lists and export datasets for downstream validation. Thus, this software passes the last hurdle for this type of project, enabling screen-based biological analysis for any interested user.

